# *Mycoplasma penetrans* Methionyl tRNA Synthetase is an Asymmetric Dimer fused to N-terminal Ancillary Domains

**DOI:** 10.1101/2025.10.13.682103

**Authors:** Behrouz Ghazi Esfahani, Madelynn K. Bowman, Rebecca W. Alexander, M. Elizabeth Stroupe

**Affiliations:** Department of Biological Science and Institute of Molecular Biophysics, Florida State University, 91 Chieftain Way, Tallahassee, FL, 32306; Department of Chemistry and Center for Molecular Signaling, Wake Forest University, 455 Vine St., Winston-Salem, NC 27101; Department of Biomedical Sciences, College of Medicine, Florida State University, Tallahassee, FL 32306, USA

**Keywords:** Methionyl tRNA synthetase, *Mycoplasma penetrans*, cryo-EM, class V pyridoxal phosphate-dependent aminotransferase, nucleotidyl transferase

## Abstract

Diverse aminoacyl-tRNA synthetase (AARS) gene fusions are now recognized as a common mechanism for enhancing genetic diversity across all domains of life. The *metS* gene from *Mycoplasma penetrans* is a striking example of such an evolutionary mechanism because although *M. penetrans* has a condensed genome, the *metS* gene is nearly twice the size of a typical bacterial gene encoding methionyl tRNA synthetase (MetRS). We used cryo-EM to analyze the structure of the MpMetRS gene product to show that it is the product of three distinct enzyme domains: an N-terminal nucleotidyl transferase, a dimeric alanine-glyoxylate aminotransferase, and a MetRS. Only the N-terminal domains show two-fold symmetry, and the MetRS domain is only partially resolved. Modelling the full structure shows that a conformational change must occur to accommodate a tRNA-bound MetRS domain. A further rearrangement of the catalytic domains would also be necessary to bring the active sites adjacent to one another if this unique assembly of catalytic domains functions to channel substrates to MetRS.

## Introduction

*Mycoplasma penetrans* is an obligate intracellular human pathogen that typically infects the respiratory or urogenital tracts of HIV-infected individuals^1^. Mycoplasma genomes are considerably smaller than in other bacterial pathogens, challenging therapeutic targeting of canonical antibiotic pathways. Specifically, the *M. penetrans* genome contains open reading frames for 1038 proteins, compared to 4288 in *Escherichia coli*, so it relies on its host for many of its basic housekeeping and metabolic needs^2^.

Despite *M. penetrans*’ minimal genome, the *metS* gene that encodes methionyl-tRNA synthetase (MpMetRS) is nearly twice the size of most bacterial MetRS enzymes^3^. Sequence homology predicts that the resulting MetRS contains a class V pyridoxal phosphate-dependent aminotransferase domain upstream of the tRNA aminoacylation domain that most closely aligns with alanine-glyoxylate aminotransferase (AGAT)^3^. In addition to the AGAT and MetRS domains, there is an N-terminal domain that harbors ambiguous sequence homology. Together, MpMetRS has the following domain organization: domain of unknown function – AGAT – MetRS. To date, no other organism with a similar chimeric MetRS has been identified, thus MpMetRS presents a unique drug target.

Many aminoacyl-tRNA synthetases (AARSs) have acquired polypeptide domains that extend function beyond the canonical tRNA aminoacylation activity^4^. These additional functions include transcriptional regulation, histidine biosynthesis, DNA binding, and mitochondrial RNA splicing^5^. Some MetRS enzymes contain a C-terminal domain that promotes dimerization and also tRNA binding^6^. Eukaryotic MetRSs are typically present in the multisynthetase complex (MSC); noncatalytic domains appended to MetRS and other MSC components drive this assembly^7^.

The MetRS and AGAT domains each perform their predicted enzyme function when expressed as full-length enzymes or when the domains are expressed in isolation: the AGAT domain catalyzes the transamination of 2-keto-4-methylthiobutytrate (KMTB) to methionine (Met) and the MetRS domain aminoacylates tRNA^Met 3^. Further, Met synthesized from KMBT by the AGAT domain can be attached to the 3’-end of tRNA^Met^ by the MetRS domain in the absence of exogenous Met^3^. How those domains are situated is unknown because there is no known structure of this unique chimeric enzyme.

Here, we report the 3.66 Å-resolution structure of MpMetRS that reveals an N-terminal nucleotidyl transferase (NTase) domain fused to a dimeric AGAT domain followed by a single helical bundle from one of the two synthetase domains. From the structure and modeled N-terminal-most domain, the 3D homology was identified as most closely aligned with an NTase domain. Further, our structure shows that the two enzyme modules organize around the AGAT-dominated two-fold symmetry axis independently such that only the N-terminal and AGAT domains follow C2 symmetry.

## Materials and Methods

### Preparation of Biological Specimens

Full length *metS* harboring a mutated codon for the M568A alteration to remove an internal start codon was cloned into pQE70 behind an N-terminal six-histidine tag. The resulting plasmid was transformed into XL-10 Gold *E. coli* (Agilent, Santa Clara, CA, USA). Cells were grown at 37 ^o^C in LB broth with ampicillin selection until O. D._600_ = 0.6, the temperature was dropped to 25 ^o^C, and then induced with 1 mM IPTG. Cells were further grown overnight at 25 ^o^C before being harvested by centrifugation at 4,000 x g for 30 min. Cells were resuspended in the working buffer: 50 mM HEPES/pH 6.8, 150 mM KCl, and 10 mM imidazole.

For purification, protease inhibitors (Roche Diagnostics, USA), 1 mM PMSF, and 1 mM pepstatin-A were added, followed by lysis with a microfluidizer and clarification by centrifugation at 13,000 x g for 30 min. Supernatant was then applied to an Ni-NTA affinity column (Cytiva, Marlborough, MA, USA) in the lysis buffer, washed with 50 mM HEPES/6.8, 500 mM KCl, and 10 mM imidazole to remove contaminating nucleic acid. After re-equilibration in the loading buffer, the sample was eluted with 250 mM imidazole and dialyzed into 50 mM HEPES/6.8, 150 mM KCl, and 10 mM imidazole. Next, the sample was loaded onto a heparin affinity column (Cytiva) to remove any residual nucleic acid-bound protein. Gradient elution to 500 mM KCl, where MpMetRS eluted at about 200 mM KCl, resulted in 99% pure full-length protein, based on SDS-PAGE analysis, with an A260:280 ratio of 0.5. Sample was polished over a Sephacryl S300 (Cytiva) size exclusion chromatography column and then concentrated to ∼10 mg/mL.

Mycoplasma penetrans tRNA^Met^ (MptRNA^Met^) was prepared by *in vitro* transcription from overlapping oligonucleotides as described in Sherlin and coworkers^8^ using sequence information from the Genomic tRNA database^9^. Transcript was purified by 8M Urea-PAGE followed by elution in 0.5 M ammonium acetate (pH 5.3), 1 mM EDTA and ethanol precipitation. tRNA was resuspended in TE buffer at −20 °C; concentration was determined by absorbance at 260 nm.

### Cryo-EM sample preparation

MpMetRS (3 μL of 0.04 mg/mL or 0.04 mg/mL MpMetRS with 5-fold excess folded tRNA^Met^) was applied to a hydrophilized graphene-coated^10^ holey carbon-on-copper Quantifoil cryo-EM grid (Quantifoil, Jena, Germany). Plunging into liquified ethane after a blot force of 4, 1s blot was performed on a Vitrobot Mark IV (Thermo Scientific, Waltham, MA, USA), resulting in thin ice and an even distribution of monodispersed particles. Alternatively, 10 mg/mL MpMetRS was applied to a nanowire self-wicking grid using the chameleon® (SPT Labtech, Melbourne, UK) nanospray device^11^ with a 54 ms wicking time.

### Cryo-EM data collection and processing

Images were collected on a Titan Krios transmission electron microscope (Thermo Fisher Scientific, Waltham, MA, USA) operating at 300 kV on a K3 camera (Gatan, Waltham, MA, USA) with the Leginon automated data acquisition package^12^, housed in the FSU-BSIR (Florida State University Biological Science Imaging Resource). For apo-MpMetRS, a total of 4,000 movies were collected with a pixel size of 0.86 Å/pixel. For MpMetRS with tRNA^Met^, 4,400 movies were collected at 1.12 Å/pixel. After motion correction using MotionCor2^13^ in the Relion-3 GUI^14^, CTF estimation was performed in CTFFIND4^15^ for both datasets. Particles were initially selected using the “blob picker” algorithm, analyzed with 2D classification to identify appropriate templates, then refined with the “template picker” in CryoSPARC^16^. All subsequent analysis was performed in CryoSPARC.

Further refinement of the apo-MpMetRS dataset by 2D classification identified 202,593 particles that were used for initial model building. Non-uniform refinement with C2 symmetry in 3D resulted in a 3.27 Å-resolution map. The map showed no evidence of the AARS catalytic domain. Closer inspection of the 2D classes identified a subset that revealed ill-defined additional features. This limited subset of particles (60,000) was used to refine a non-uniform, asymmetric map to 3.66 Å resolution. All maps were sharpened with deepEMhancer for enhanced visualization^17^.

For the tRNA^Met^/MpMetRS specimen, 424,722 particles were used for 2D analysis and refined using the same protocol. Further 3D analysis revealed a similar structure of equivalent resolution to that of apo MpMetRS.

### Model building and refinement

Model building was initiated with the ModelAngelo^18^ model builder. Iterative real-space refinement in PHENIX^19^, with manual fitting in Coot^20^, was performed independently for the core dimer and the C-terminal domain helical bundle of the AARS domain.

After model building of the N-terminal most domain, the atomic coordinates were submitted to the DALI server^21^. Out of the first 34 matched results with z-score ≥10, 26 (76%) are nucleotide transferase domains. The highest z-score belongs to a Nucleotidyl Transferase/Aminotransferase, Class V (6PD1-B^22^).

## Results

### The AGAT domain forms a canonical dimer fused to an N-terminal domain of unknown function

Initial 2D analysis shows classes with high-resolution features that correspond to the dimer mediated by the AGAT domain (Fig. 1A). Refinement and 3D analysis of all 202,593 particles further supports this observation, showing only density for the dimeric core that resolves to 3.27 Å resolution (Fig. 1B).

**Figure 1:**
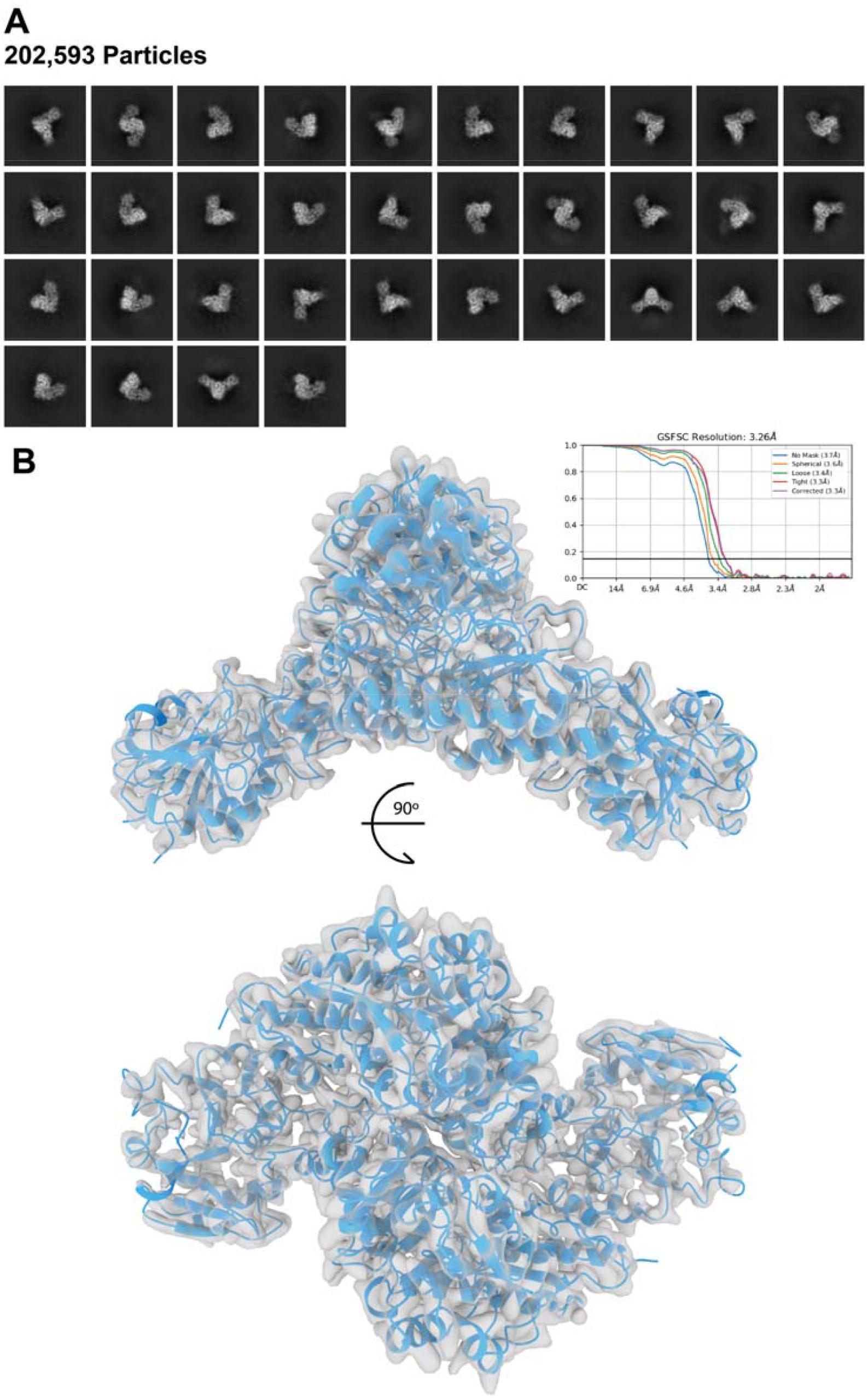
**A)** Selected 2D classes with high resolution features. **B)** 3.27 Å resolution structure from the particles used to calculate the 2D classes in (A), used to model the core dimer including the AGAT domain and the N-terminal unknown region.

A *de novo* model of the AGAT domain shows that the interface is mediated by salt bridges, hydrophobic stacking interactions, and hydrogen bonds, akin to other dimeric amino transferases like the alanine-glyoxylate aminotransferase from *Anabaena* 7120^23^ (Figs. 2 and S1), which is 27% identical and 45% similar. For example, the salt bridge between Arg297 and Asp419 is strictly conserved between the two. A stacking interaction between Ile294 and Lys416 in the MpMetRS AGAT domain is analogous to a stacking interaction in the *Anabaena* AGAT between Met155 and Trp240. The *en face* interaction between Pro209 and Thr436 is also conserved (Pro30/Thr260 in the *Anabaena* homolog).

**Figure 2:**
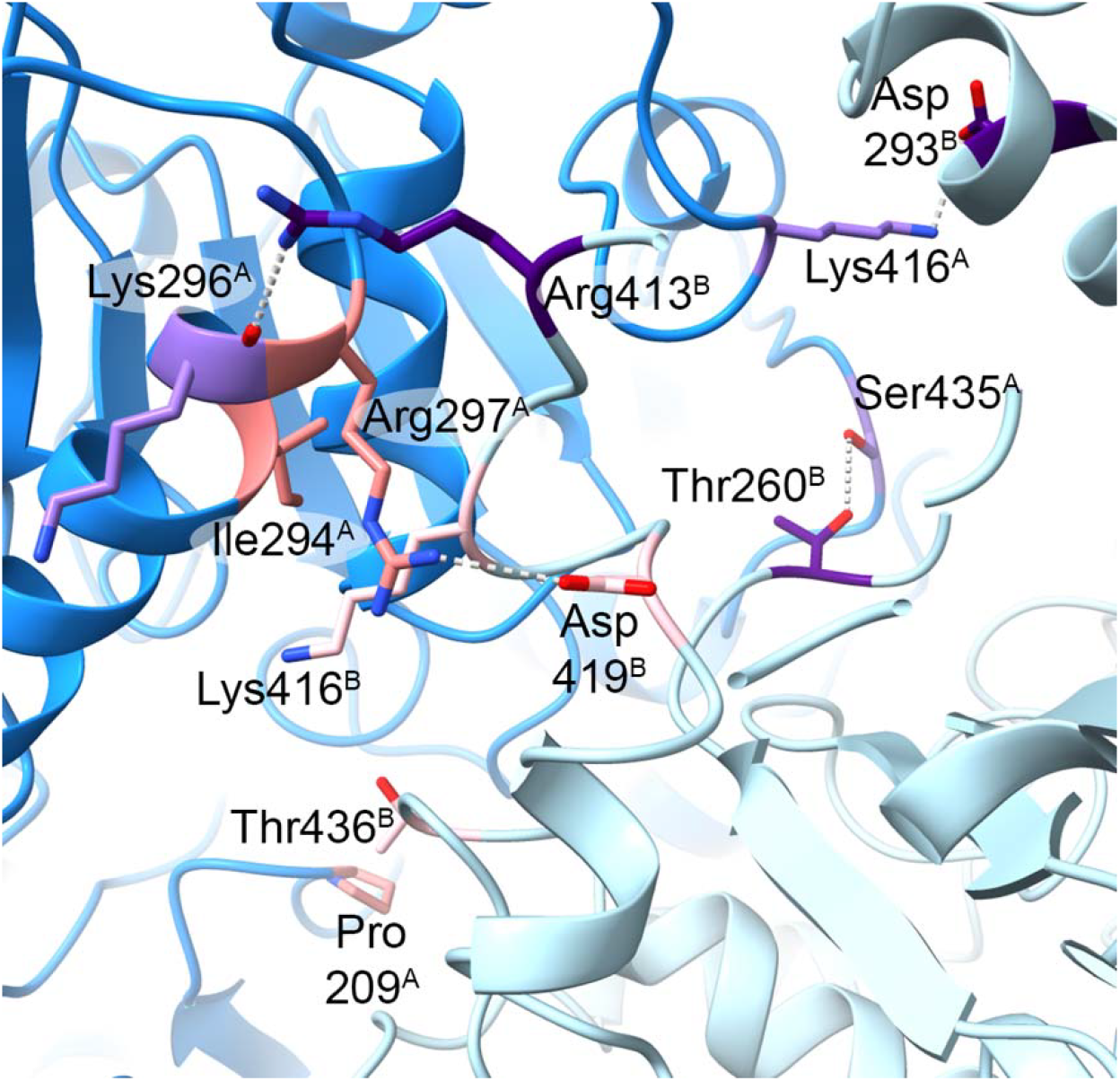
The dimeric interface for the AGAT domain of MpMetRS, modeled from the cryo-EM density, has some conserved (light/dark pink) and some unique (light/dark purple) interactions.

In other instances, the MpMetRS interface harbors unique interactions despite some sequence conservation (Figs. 2 and S1). For example, Arg413, which is conserved with *Anabaena* Lys236, forms a hydrogen bond with the carbonyl from Lys296. In the *Anabaena* homolog, the homologous Lys236 forms a salt bridge with the carboxylate from Asp114. The homologous Asp in MpMetRS (Asp293) also participates in a salt bridge with a lysine, in this case the non-conserved Lys416. In MpMetRS, Thr260 forms a hydrogen bond with Ser435. Conserved Thr83 from *Anabaena*, on the other hand, rotates to interact with the phosphate group from the PLP cofactor.

Despite the canonical aminotransferase dimer, its position between a novel N-terminal domain and a C-terminal AARS is unique to *M. penetrans*, as evidenced by sequence analysis of the chimeric MetRS^3^. Each of the three domains folds independently, but only the N-terminal-most and AGAT domains exhibit C2 symmetry, described in detail below (Fig. 3). Structural homology of the N-terminal-most domain reveals that it is similar to NTase enzymes, with the highest similarity to the N-terminal domain of GlmU from *Mycobacterium tuberculosis*^24^. In contrast to the large interface between the AGAT domains, the NTase domain is at the periphery of the dimer and tucks into a pocket formed at the AGAT dimer interface (Fig. 3). The predicted AGAT active site sits about 40 Å away from the predicted nucleotide binding pocket on the NTase domain from the same polypeptide and about 50 Å away from the nucleotide binding pocket of its partner.

**Figure 3:**
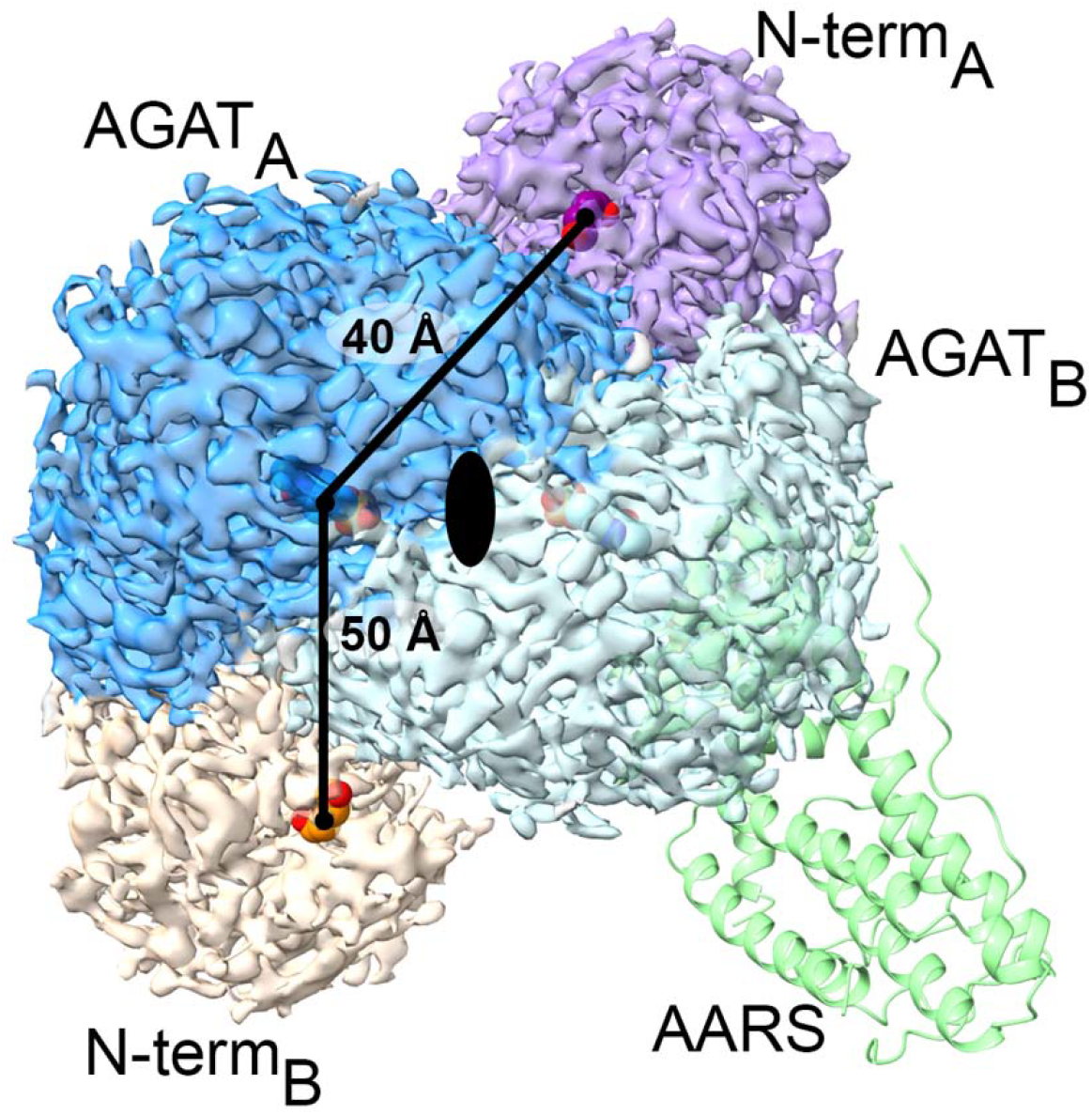
The N-terminal-most domain of MpMetRS (purple and peach) tucks into a pocket formed by the interface between the intermediate AGAT dimer (dark and light blue). The approximate active sites for the N-terminal NTase and AGAT domains were determined by superimposition on their most closely aligned homologs with bound ligand and represented as space-filling spheres (PDB 6GE9, GlmU^24^, for the NTase domain and PDB 1VJO, alanine-glyoxylate aminotransferase^23^, for the AGAT domain.

### The position of the AARS domain does not follow C2 symmetry

To better understand the relationship between the NTase, AGAT, and AARS domains, we performed further 3D analysis of a subset of particles that exhibit additional, less-well defined features in 2D (Fig. 4A). The resulting structure shows additional density with helical features (Fig. 4B). Modeling of the AARS domain confirms homology to other similar class Ia AARSs where the C-terminal anticodon-binding domain forms an antiparallel five-helical bundle. There is no well-resolved density for the N-terminal catalytic site of the AARS or for the AARS from the opposing subunit, despite PAGE-analysis showing the purified full-length protein that is predominantly a dimer (Fig. S2).

**Figure 4:**
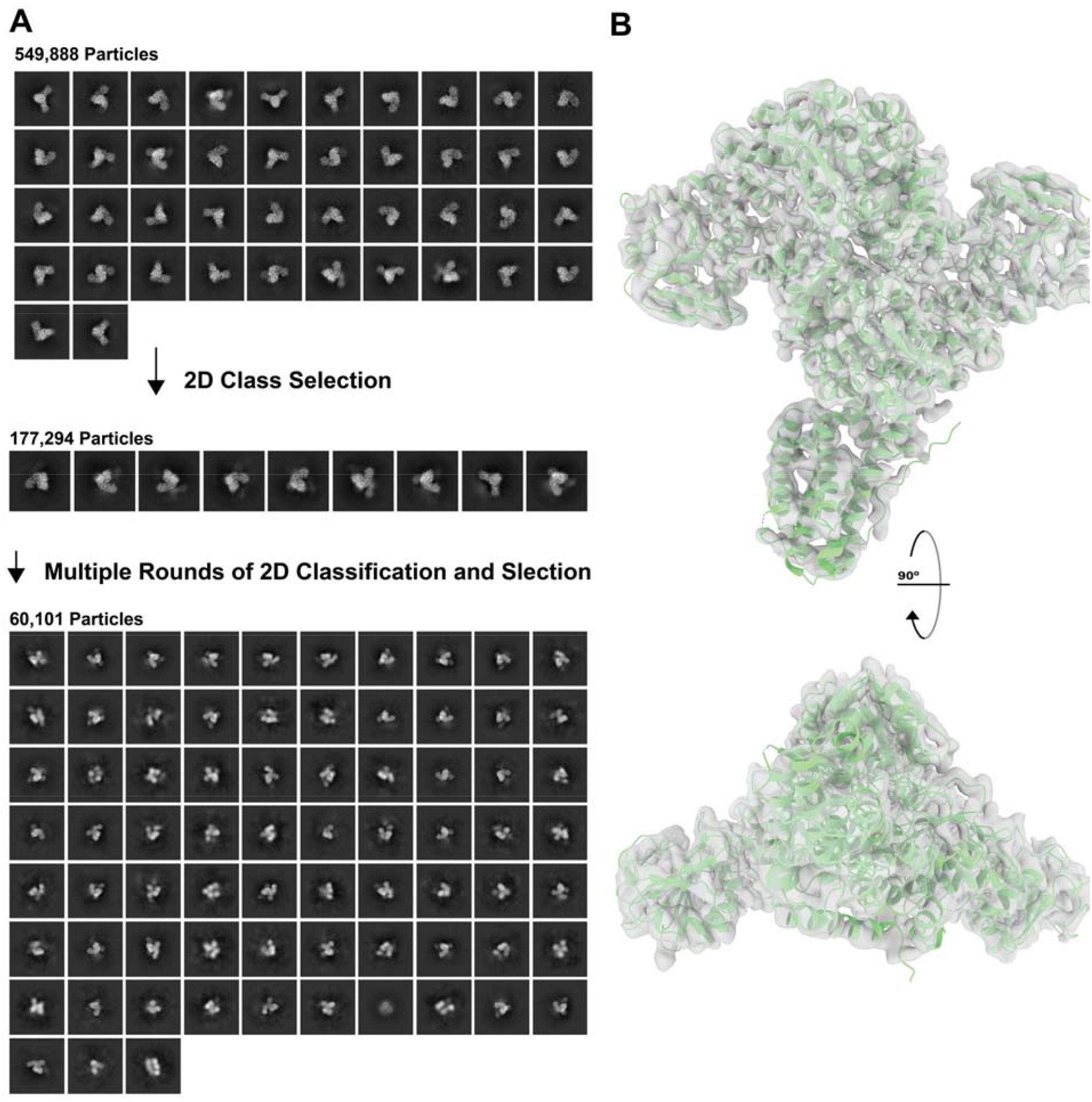
**A)** Further 2D classification of the apo MpMetRS particles from the classes with faded features results in a set of classes with additional features. **B)** EM map from the particles selected at (A) to model the helical bundle from the C-terminal catalytic domain.

In contrast, when sample was pre-mixed with excess tRNA, additional 2D classes appeared with a higher proportion of particles (40% compared to 0% without tRNA) that showed ill-defined density at the outside of the central, well-defined dimer (Fig. S3). Despite extensive 3D classification, however, no high-resolution features appeared for the catalytic domain or bound tRNA.

### MpMetRS oligomeric state depends on the protein concentration

The above results were determined from a 0.04 mg/mL sample applied to graphene-coated grids. In contrast, the chameleon® plunging system (SPT Labtech) allows for sample application at a higher concentration^11^. 2D analysis of this preparation shows additional classes that appear with additional, well-resolved densities (Fig. 5A). Further 3D analysis of these subsets of particles reveals tetrameric and hexameric NTase-AGAT assemblies (Fig. 5B). Within those higher-order oligomers, the NTase and AGAT domains appear concatenated, with the NTase domain of one polypeptide positioned in opposition to the AGAT domain of the next polypeptide. No density for the AARS domain was observed in these particles.

**Figure 5:**
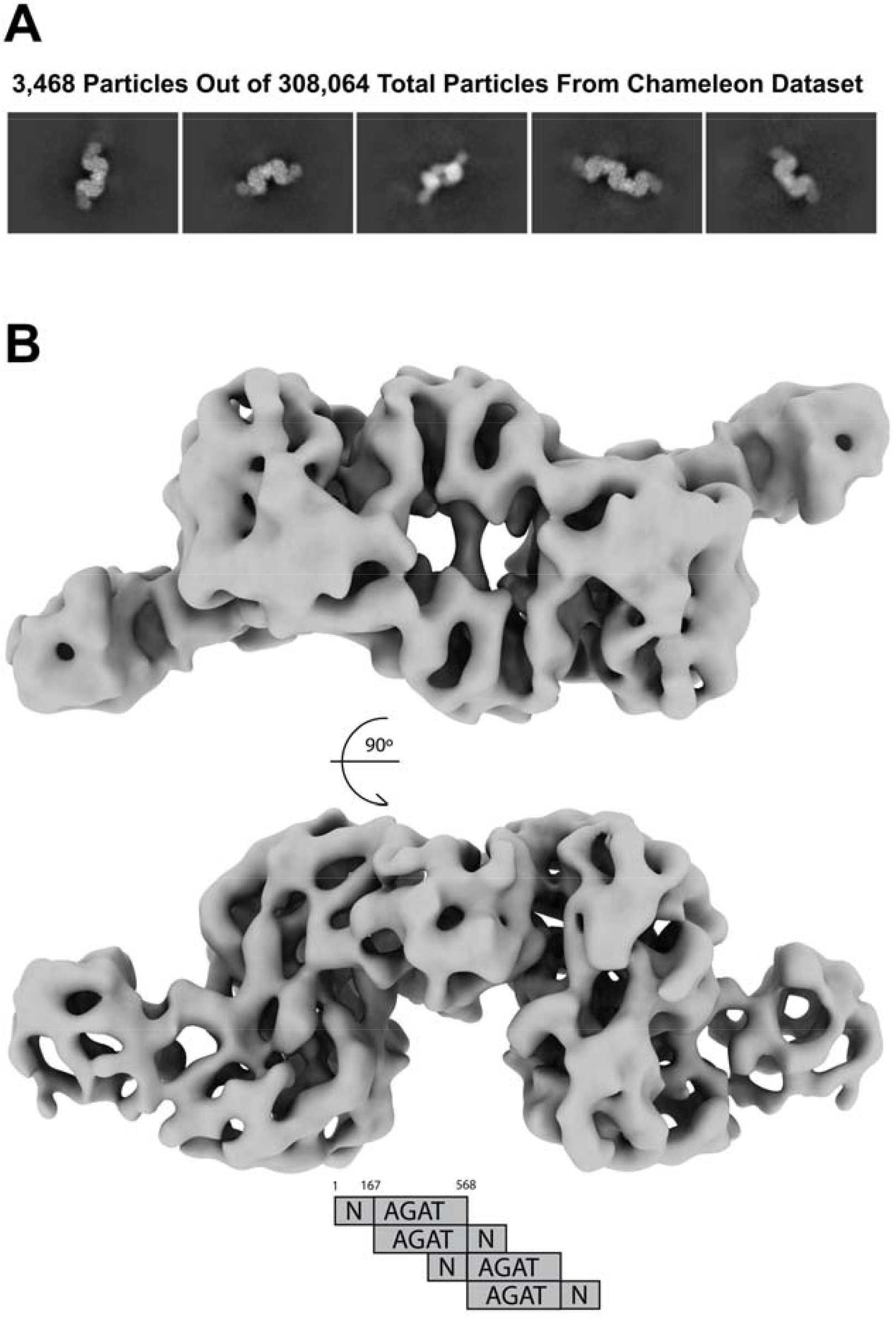
**A)** MpMetRS 2D classes that show higher-order oligomerization, obtained from particles vitrified with the chameleon ^11^. **B)** Low resolution 3D reconstruction of a tetrameric MpMetRS that shows density only for the N-terminal and AGAT domains, obtained from the subset of particles shown in (A).

### Modelling

The loop that connects helix 2 and 3 of the AARS helical bundle tucks into the AGAT domain in a crevice formed between two of its helices at the opposite face of the dimer interface (Fig. 6A). Docking the helical bundle from monomeric *E. coli* MetRS^25^ predicts that the catalytic domain would fall immediately between the C2-symmetric NTase domains (Fig. 6B). In contrast, docking the tRNA-bound *E. coli* CysRS^26^ onto the *M. penetrans* helical bundle shows that the tRNA would sterically clash with the N-terminal chimera as positioned in the apo structure (Fig. 6C).

**Figure 6:**
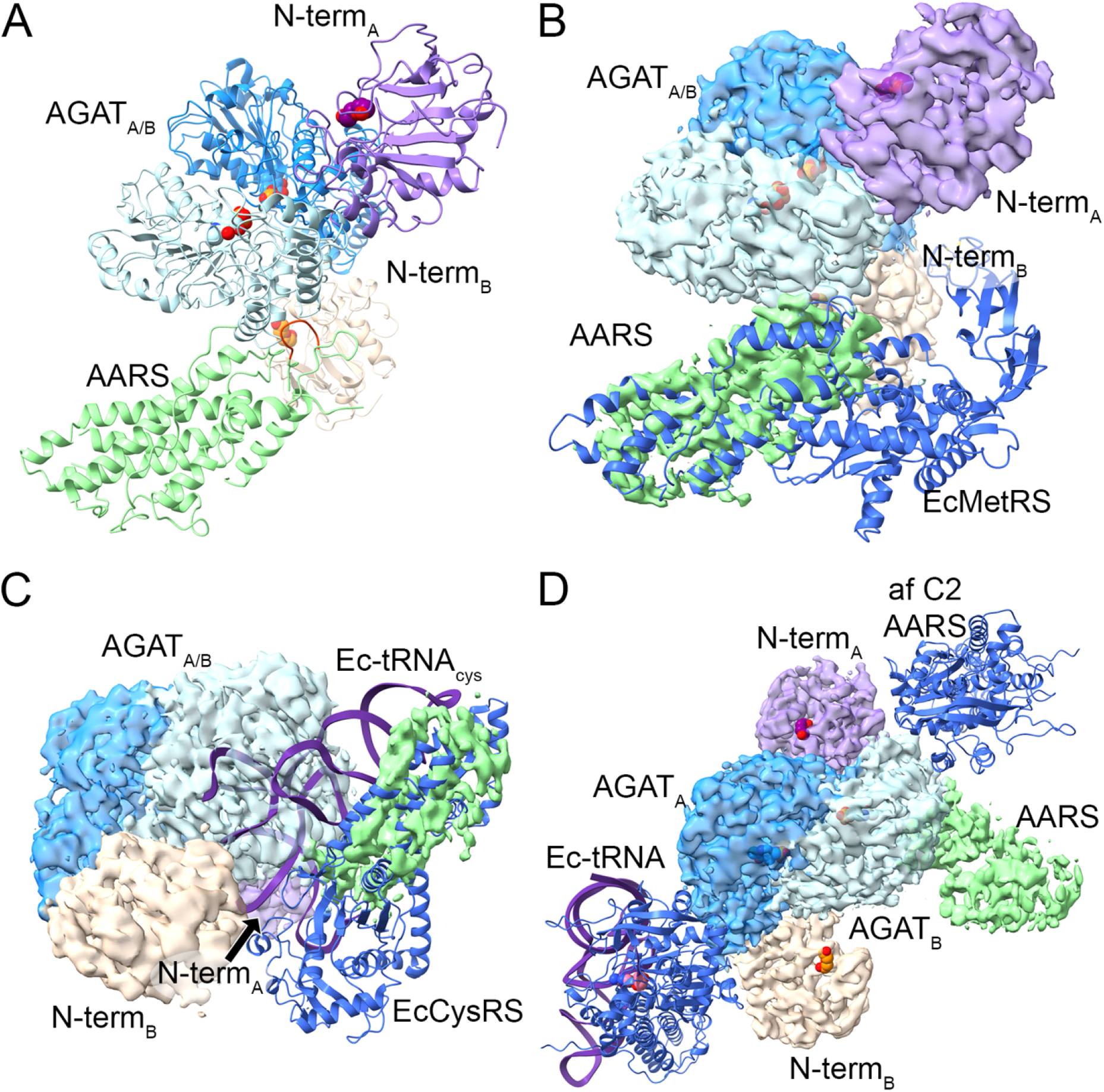
**A)** The helical bundle from the tRNA anticodon recognition domain (green) binds through a helix-loop-helix (red) motif between two helices from the AGAT domain. **B)** Docking the full MetRS structure from *E. coli* ^25^ (blue) onto the helical domain from MpMetRS places the acceptor stem in the cleft between MpMetRS’s C2-symmetric N-terminal NTase binding domains (peach and purple). **C)** Docking the tRNA-bound CysRS from *E. coli* ^26^ in a similar fashion identifies potential steric overlap between tRNA binding and the positioning of the AARS relative to the AGAT dimer. **D)** AlphaFold2^27^ modelling of the fully C2-symmetric MpMetRS suggests the tRNA could bind at the extreme periphery of the dimer with an approximately 60° rotation of the AARS domain.

To better understand how this steric clash could resolve, we compared the experimental results, which show an asymmetric and incompletely ordered dimer, with those from AlphaFold2^27^, imposing C2 symmetry. To adopt the fully symmetric dimer, the helical subdomain from the AARS domain would need to rotate about 60° using the AGAT active site as a pivot point reference (Fig. 6D). With that rotation, the AARS catalytic domain is removed from the space between the two N-terminal NTase domains, opening the tRNA binding site. The 3’ end of the tRNA sits about 46 Å away from the AGAT active site.

## Discussion

The MpMetRS protein is included amongst a now growing class of AARS enzymes where the aminoacylation domain is expressed as a chimera with a “helper” protein, often of housekeeping function^28^. The triple fusion of a NTase, AGAT, and AARS is, to date, unique. *M. penetrans* has a minimal genome, relying on the human host for many metabolic and regulatory functions. In this way, the structural relationship of those domains furthers our understanding of why this assembly was evolutionarily favorable.

Structural, not sequence, homology of the N-terminal-most domain identifies it as most closely resembling the Rossmann fold dinucleotide binding motif from MtGlmU. In the case of MtGlmU, the NTase activity is responsible for adding a uridylyl group to the UDP-*N*-acetylglucosamine precursor to peptidoglycan^24^. Interestingly, MtGlmU is also the product of a gene fusion event with the enzyme that catalyzes the prior step in peptidoglycan synthesis, the acetyl-CoA dependent acetylation of glucosamine 1-phosphate. Optimal activity of both sites depends on the protein’s trimeric assembly^29^, suggesting precedence for the functional relevance of chimeric nucleotide-dependent enzymes. In the case of MpMetRS, perhaps the addition of an NTase domain increases the local concentration of ATP to catalyze the addition of the methionyl group to charge the tRNA.

To further explore this idea, we modeled the apo anticodon binding subdomain of the AARS domain based on the crystal structure of EcMetRS. This simple docking positions that subdomain between two NTase domains, but on the opposite face from the putative NTase active site. When the same docking is performed using a tRNA-bound AARS, EcCysRS^26^, a conflict arises from the tRNA binding pose in relationship to the cleft between the central, C2-symmetric N-terminal NTase/AGAT. Likely a large conformational change occurs, perhaps through the linker between the core AGAT and the AARS domains, to perform the aminoacylation of Met-tRNA. This linker region is disordered in apo MpMetRS such that it is not visible in the 3D reconstruction, despite extensive classification. For that reason, we cannot assess whether the visible helical domain, which is C-terminal to the aminoacylation sub-domain, originates from chain A or chain B.

The zinc-binding motif subdomain from the AARS putatively projects into the space between the N-terminal NTase and the AGAT of a continuous polypeptide. Molecular dynamics of the highly similar EcMetRS, which lacks the N-terminal additional domains, predicts that this region is particularly mobile^30^. The space between the C2-symmetric NTase domains accommodates this domain, which is sufficiently mobile in the apo form of the enzyme that it is not visible in the reconstruction even in the absence of imposed C2 symmetry. This conformation of the AARS domain positions the tRNA binding interface against the central dimer, however, such that there would be significant steric clashing if the tRNA were to bind with the AARS as it is in the apo structure. Specifically, the AARS helix-turn-helix motif that interacts with the AGAT domain binds to the inside elbow of the tRNA when the substrate is bound, thus the apo conformation of the AARS domain relative to the AGAT domain is in conflict with the substrate-bound conformation.

*E. coli* MetRS (EcMetRS) is dimeric *in vivo*, but a monomeric construct expressed without the C-terminal dimerization domain is functional *in vitro*. Prior molecular dynamics (MD) simulations of EcMetRS were performed using coordinates from the high-resolution crystal structure of monomeric MetRS^25^, which shares 25% sequence identity and 45% sequence similarity with the AARS domain of MpMetRS^30^. Several regions of the protein exhibited high mobility, most notably a zinc-binding domain inserted between halves of the enzyme active site, a surface loop that serves as a cap to the active site, and the anticodon binding domain. Each of these three regions has been implicated in MetRS function, suggesting that protein dynamics contribute to activity. The MD simulation was carried out on the ligand-free protein; it is certainly anticipated that tRNA binding would alter protein dynamics.

Oligomerization of nucleotide and polynucleotide binding proteins is not atypical, for example in the DNA recombination protein RecA^31^, cytidine triphosphate synthase^32^ or the G-quadruplex binding protein nucleoside diphosphate kinase^33^. We see this at high concentrations of MpMetRS, such that the N-terminal NTase domains dimerize orthogonally to the AGAT dimeric interface through an active site loop (Figs. 5 and 7). A secondary interaction between an NTase and an AGAT domain from a separate polypeptide further forms while maintaining an open channel to the NTase active site (Fig. 7, inset), so presumably the substrate and product can still diffuse into and out of the binding pocket. Although the AARS domain is not sufficiently ordered to see its density, there is no steric conflict between its position in the dimer and the octamer.

**Figure 7:**
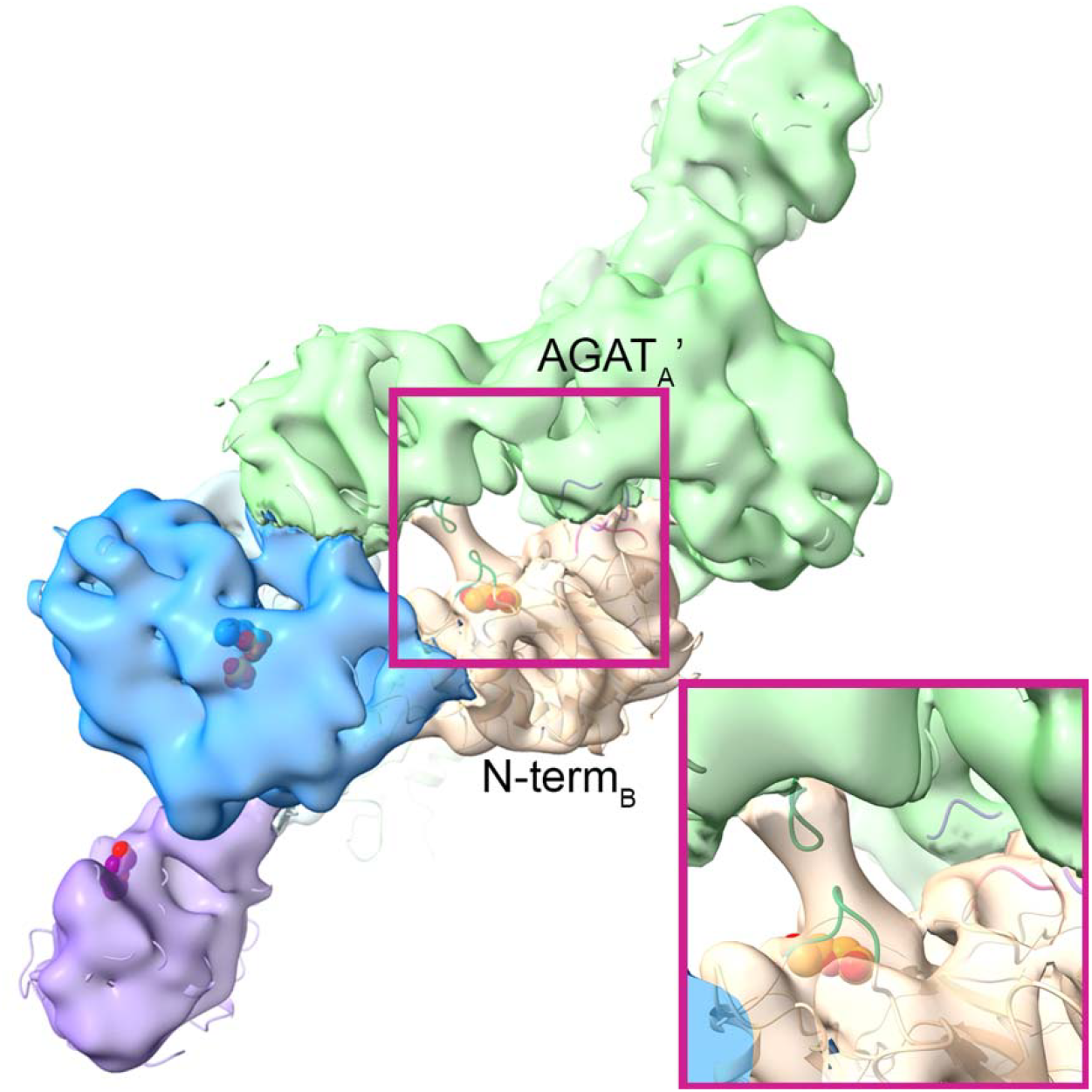
Concatenation of the N-terminal and AGAT domains forms a secondary two-fold symmetry axis between N-terminal NTase domains (peach and green, active site loop in dark green) and a new interface between the AGAT domain (green or blue, involved loops in purple) and an opposing NTase domain (peach or green, involved loop in dark pink). There is no density for the AARS domain.

### Conclusions

Coordination of the three activities might explain the evolutionary advantage of having each activity co-localized within a single polypeptide, even in an organism that has a minimal genome. First, the NTase domain increases the availability of the precious cellular resource ATP. Second, the AGAT domain synthesizes methionine to supply the aminoacylation catalytic domain. Finally, the mobile AARS domain positions its catalytic site to receive these essential metabolites before binding tRNA^Met^.

## Data availability

The refined coordinates were deposited in the Protein Data Bank as 9OS7. The cryo-EM map and half maps were deposited in the Electron Microscopy Data Bank under the code EMD-70794.

## Acknowledgements

We thank Dr. Christopher Stroupe and Jen Kennedy for helpful discussions. Florida State University supports cryo-EM in the Biological Imaging Resource Center, which houses the following equipment used in this study: a Gatan Solaris Plasma Cleaner (NIH grant S10 RR024564), a Hitachi HT7800 (NSF grant MRI2017869 to M.E.S.), a ThermoFisher Vitrobot Mark IV (NIH grant S10 RR024564), an SPI chameleon® plunging system (NIH grant R24 GM145964), a ThermoFisher Titan Krios (NIH grant S10 RR025080), and a DE Apollo direct electron detector (NIH grant R35 GM139616). This work was further supported by National Science Foundation grants MCB1856502 and CHE1904612 to M.E.S.

