## Supplemental Material for "*Mycoplasma penetrans* Methionyl tRNA Synthetase is an Asymmetric Dimer fused to N-terminal Ancillary Domains"

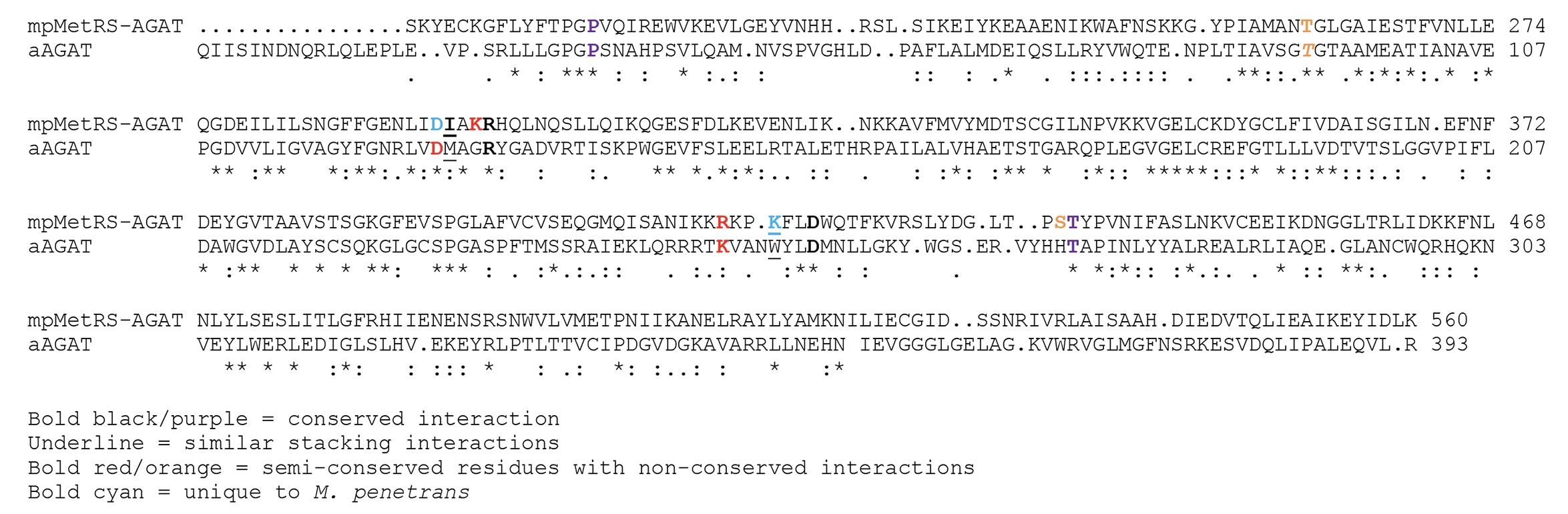

Figure S1. Structure-based sequence alignment of the *MpMetRS* AGAT domain with the AGAT from *Anabaena* (PDB 1VJO^1^)

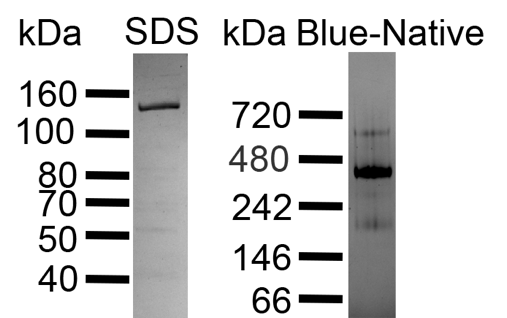

Figure S2. SDS PAGE gel (left) and blue Native-PAGE gel (right) showing the purity and oligomeric state of MpMetRS.

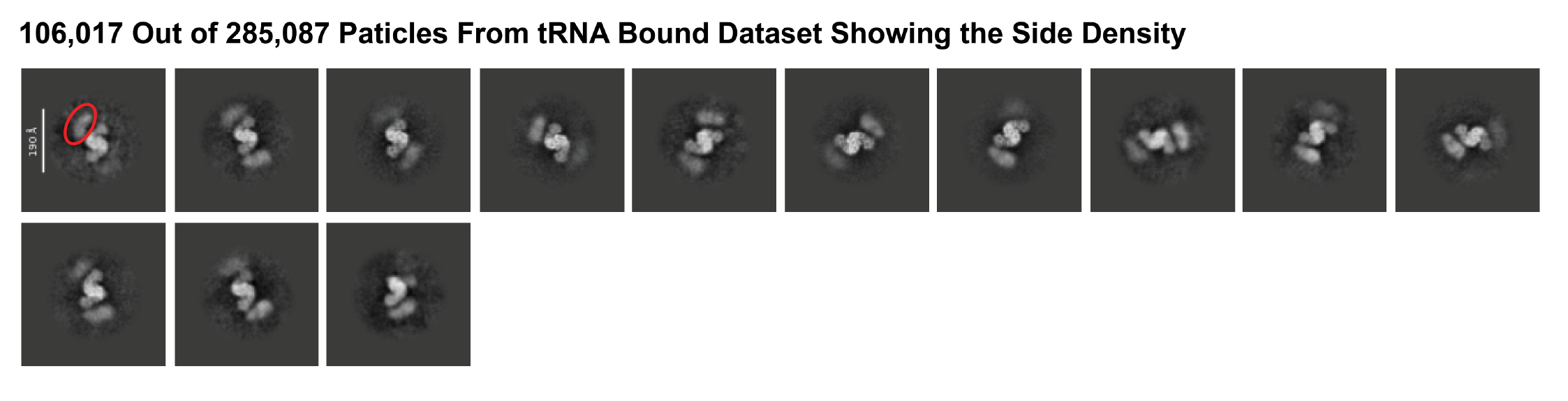
Figure S3. A subset of particles from the tRNA bound dataset showing the peripheral side density, for example marked by a red oval.
